# A slippery slope: assessing the amphibian extinction crisis through the lens of climate refugia

**DOI:** 10.1101/2024.09.25.615003

**Authors:** Desiree Andersen, Amaël Borzée, Yikweon Jang

**Affiliations:** School of Natural Resources and the Environment, University of Arizona, Tucson, USA; Division of EcoScience, Ewha Womans University, Seoul, Republic of Korea; Laboratory of Animal Behaviour and Conservation, College of Life Sciences, Nanjing Forestry University, Nanjing, China

## Abstract

In a time of increasing climate uncertainty, it is ever important to identify and preserve refuges for species, optimizing the effort to cover the largest possible number of clades. Climatic refugia are areas of climatic stability that will remain suitable for species over time under multiple climate change scenarios. Amphibians are important indicators of biodiversity and ecosystem health, in addition to being the most threatened class of vertebrates and can therefore serve as proxies to designate protected areas. In this study, we use ecological modeling to delineate current distributions of all amphibian species with adequate data globally. We then projected these distributions to four climate change scenarios, and four time periods, under two climate models representing low and high climate sensitivity. Climate refugia by species were calculated as the average of all climate change scenarios and time periods and were calculated separately for the two climate models of low and high sensitivity. We additionally extracted areas of current distributions and refugia to geographic regions, Udvardy biomes, and global protected areas. With the high climate sensitivity model (CNRM-CM6-1), 2,959 species would experience a refugia area that is reduced from their current distribution, with 139 species having no refugia area. Under the low climate sensitivity model (MIROC6), 2,864 species would have a reduction in area, with 136 having no refugia. While both climate models yielded similar results in terms of percent change, the MIROC6 model overall had less of a negative impact on species’ refugia. The results of this study present a somber warning of the amphibian extinction crisis in contrast to some of the recent literature, as well as encouragement for managers to act in order to preserve species and the ecosystems they represent.

## Introduction

Climate change has become a critical issue affecting the survival and distribution of many species worldwide, and therefore closely linked to the ongoing biodiversity crisis. The decline in species diversity poses significant challenges for maintaining the resilience of ecosystems in terms of ecological services provided by species for the continuity of ecosystems. The effects of climate change on biodiversity are expected to intensify in the coming decades, exacerbated by other anthropogenic influences including habitat loss and fragmentation, invasive species, and diseases. Among the main threats to biodiversity posed by climate change are shifts in species distributions (Borzée et al., 2019; Loarie et al., 2009; Moritz et al., 2008; Parmesan and Yohe, 2003; Wilson et al., 2005), changes in phenology (Blaustein et al., 2001; Forchhammer et al., 1998; Socolar et al., 2017), and altered food webs (Albouy et al., 2014; Rosenblatt and Schmitz, 2016; Zhang et al., 2017). Moreover, climate induced range shifts that are expected to push species towards higher latitudes and elevations (Houghton, 1996; Walther et al., 2002) are also expected to impact amphibians (Pounds et al., 1999) which are more vulnerable to displacement as they do not disperse over long distances (Smith and Green, 2005).

Climate change is particularly relevant in the case of amphibians, which are among the most sensitive and vulnerable groups of animals to environmental changes (Luedtke et al., 2023). Amphibians are particularly vulnerable to these changes due to their dependence on both aquatic and terrestrial habitats, sensitivity to temperature fluctuations, and permeable skin that makes them more susceptible to changes in moisture levels and environmental toxins (Mann et al., 2009). These sensitivities and their diverse but weakly overlapping range of ecological niches make amphibians ideal biodiversity and environmental indicators of the health of both aquatic and terrestrial ecosystems (Blaustein et al., 2010; Caruso et al., 2014; Connette et al., 2015).. Therefore, amphibians are a key group for studying the impacts of climate change and monitoring the state of the environment.

Amphibians are declining at a much higher rate than other taxa, with many of the factors contributing to their decline linked to human activities, which include additive threats of habitat loss, pollution, fragmentation, and degradation, as well as the introduction of invasive species and emerging diseases (Hof et al., 2011; Luedtke et al., 2023). Amphibians have experienced substantial population declines worldwide, with 44.3% of species experiencing decreasing populations, 25.1% stable or increasing and 30.1% unknown (IUCN, 2023). Many species are now threatened with extinction as 40.2% are assessed as critically endangered, endangered, vulnerable or near threatened. Further, 15.3% of species are categorized as data deficient, making the number of threatened species underestimated.

Compounded with climate change and habitat loss, invasive amphibian species and amphibian diseases are increasingly pressing threats to the survival of amphibians worldwide. Invasive species can outcompete and prey upon native species, and have even been implicated as drivers in up to 31% of amphibian extinctions globally (Blackburn et al., 2019). Amphibian diseases, such as chytridiomycosis, have been responsible for significant declines in many species of amphibians worldwide (Fisher et al., 2021), leading to the extinction of 90 species worldwide (Scheele et al., 2019). The spread of these diseases is facilitated by global trade and transport of wildlife, and it is often challenging to control and eradicate diseases once introduced into a population.

Climatic stability refers to areas that remain climatically stable during periods of environmental change across a range of climate change scenarios and time periods (Loera et al., 2017; Yannic et al., 2014). Areas of high climatic stability could serve as refuges for species facing climate-driven habitat suitability shifts (Gavin et al., 2014; Olson et al., 2012). These areas, known as refugia, can be critical for the survival of species in the face of rapid climate change. With the ongoing effects of climate change, identifying these areas of stability will be a tool for developing effective conservation strategies that protect not only individual species but also the ecosystems and services they provide.

In the current study, we apply fine-tuned ecological modeling (MaxEnt; Phillips et al., 2017) to identify global regions of amphibian richness and predicted climate refugia across climate change scenarios for two climate models with low (MIROC6; Tatebe et al., 2019) and high (CNRM-CM6-1; Voldoire et al., 2019) climate sensitivity. We further analyze species’ current ranges and refugia areas extracted to geographic regions, Udvardy biomes (Udvardy and Udvardy, 1975), and global protected areas. Most importantly, we identify a minimum subset of protected areas for each amphibian order to preserve all species with current presence and predicted refugia in protected areas.

## Methods

### Occurrence data

Occurrence data was obtained from GBIF (2022) for the Class Amphibia by filtering coordinates without geospatial issues, specimen/human observation/occurrence. This process identified 5,794,955 records initially. To limit spatial autocorrelation, occurrences were reduced by species, latitude, and longitude at 0.1 decimal degree using the ‘aggregate’ function in the ‘stats’ package (Becker et al., 1988). Finally, species with five or more occurrences after this refinement were selected for modeling. This resulted in a total of 1,865,066 unique occurrences of 4,035 species.

### Ecological niche modeling

Species were modeled using maxent (maxent.jar from the ‘dismo’ package (Hijmans et al., 2017) with the ‘ENMevaluate’ function from the package ‘ENMeval’ (Kass et al., 2021) in R ver. 4.1.3. Environmental variables consisted of 19 bioclimatic variables (Worldclim version 2.1 downloaded from Worldclim.org; Fick and Hijmans, 2017) at a global extent and a 0.0417 decimal degree resolution (∼3.7km).

First, the environment was clipped to an extent extending five decimal degrees from a bounding box containing occurrence points for each species. This allowed for shorter processing times and limiting background points to accessible areas. Models used 10000 background points, as is standard for maxent, randomly created (‘randomPoints’ function in ‘dismo’ package (Hijmans et al., 2017)) within the clipped extent and where environmental data was available. For species with less than 25 occurrences, the partition setting was set to jackknife (Pearson et al., 2007; Shcheglovitova and Anderson, 2013). For species with 25 or more occurrences, the partition was set to random k fold using five folds. The beta multiplier (randomization multiplier) was set from 1 to 4 (including all integral values), and factors were set to linear (L), linear-quantile (LQ), linear-quantile-hinge (LQH), and hinge (H).

Best models for each species were selected on the corrected Akaike information criterion (AICc) wherein the best model was the model which delta AICc was equal to zero. These selected models were checked for model fit using area under the curve (AUC). Models were retained only if the AUC was over 0.7.

These best models were then projected to current and future climates consisting of: two global climate models (GCMs) representing high (CNRM-CM6-1; Voldoire et al., 2019) and low (MIROC6; Tatebe et al., 2019) sensitivity; for four Shared Socio-economic Pathways (SSPs; 126, 245, 370, 585); over four time periods (2021-2040, 2041-2060, 2061-2080, 2081-2100). The four SSPs represent: “sustainability” (SSP 1.26), “middle of the road” or “regional rivalry” (SSP 2.45), “inequality” (SSP 3.70), and “fossil-fueled development” (SSP 5.85) (Riahi et al., 2017). All future climate layers were downloaded from Worldclim.org. For the current climate, the projected models were thresholded at the maximum training sensitivity plus specificity threshold obtained from the corresponding maxent result file. These thresholded models were then converted to polygons, and only polygons containing occurrence points were dissolved using the ‘aggregate’ function. To simulate potential migration due to changing climates, projected models for future climates were masked to a buffer of the dissolved polygon, with a buffer distance corresponding to the average active dispersal distance for each amphibian order (Smith and Green, 2005), multiplied by number of years since 2020 to the midpoint of each time period range (i.e. the time period 2021-2040 would have this distance multiplied by 10, and so on).

For each of the two GCMs, all projections were averaged to create a single stability model, then thresholded at the maximum training sensitivity plus specificity threshold. These stable areas, or refugia, were then converted to polygons and projected to the NSIDC EASE-Grid 2.0 Global projection (EPSG: 6933) to determine their area in km^2^.

### Area analysis for databases

To determine current and refugia coverage by region, biome, ecoregion, existing protected areas(IUCN and UNEP-WCMC, 2022), and KBAs, we implemented the Zonal Statistics as Table tool in ArcGIS Pro 2.7.3 (ESRI, Redlands, CA, USA). We used protected area polygons for feature zone data, WDPA_PID (protected area identifier in the Protected Planet database) as the zone field and individual species presence and future refugia as input value rasters. The results of this analysis were summarized by order (Anura, Caudata, Gymnophiona). We finally identified a minimum subset of protected areas for each order to preserve all species with current and refugia presence in protected areas.

### Post-hoc analysis on area metrics

Percent change in area by species was tested in linear and analysis of variance models to determine, for each order, which variables might affect the extent to which species will be affected by climate change. For these post-hoc models, percent change in CNRM-CM6-1 and MIROC6 was predicted using family, latitudinal range of occurrence, and absolute value of average latitude of each species as predictor variables. Significant differences between families were ascertained using TukeyHSD. Finally, extracted areas by species within biomes and regions were tested for differences between the current area and CNRM-CM6-1 and MIROC6 refugia areas. This was tested using paired t-tests.

## Results

In total, 4,035 species of amphibians were selected for modeling (Table S1). Of these, there were 3,495 species of Anura, 484 species of Caudata, and 57 species of Gymnophiona. Across all models, area under the curve (AUC) and true skill statistics (TSS) values averaged 0.9420 ± 0.0510 and 0.7966 ± 0.1352, respectively, both of which are considered to represent excellent fit (Table 1). Of 50,119 protected areas considered, 49,970 had predicted suitability for at least one Anura species, 33,088 for at least one Caudata species, and 4,552 for at least one Gymnophiona species. Of these, 409 (Table S2) could be selected as a minimum subset covering 3,394 (97.1%) of Anura species currently, 3,273 (93.6%) species’ refugia in CNRM-CM6-1 and 3,287 (94.0%) species’ refugia in MIROC6. For Caudata, 157 protected areas cover 460 (95.4%) species currently, 427 (88.6%) species’ refugia in CNRM-CM6-1 and 420 (81.7%) species’ refugia in MIROC6. Finally, 27 protected areas currently cover 49 (86.0%) of Gymnophiona species, 41 (71.9%) of species’ refugia in CNRM-CM6-1 and 42 (73.7%) of species refugia in MIROC6. Due to some overlap between orders, this totals to 539 protected areas.

**Table 1:** By order, average AUC, current suitable area, and percent change in refugia from the current suitable area for MIROC6 and CNRM-CM6-1 global climate models. Parentheses represent number of species with no refugia in future climate scenarios. Models were generated using MaxEnt with model selection by ENMeval (Kass et al., 2021).

| Order | AUC | TSS | CNRM-CM6-1 $\Delta$ | MIROC6 $\Delta$ |
| --- | --- | --- | --- | --- |
| Anura | 0.9401 ± 0.0509 | 0.7892 ± 0.1346 | -3.6% (99) | -2.4% (93) |
| Caudata | 0.9565 ± 0.0492 | 0.8565 ± 0.1204 | -34.1% (32) | -29.8% (37) |
| Gymnophiona | 0.9356 ± 0.0524 | 0.7473 ± 0.1592 | -37.5% (8) | -32.6% (6) |

On average, stable area (above TSS threshold for average of all model projections) by species for the CNRM-CM6-1 climate model was reduced from the current suitable area by 7.7%, while the stable area for the MIROC6 climate model was reduced by 6.1% (Figure 1; see Table 1 for changes by order). For the CNRM-CM6-1 climate model, 2,959 species experienced a reduction of stable area from current suitable area, while 1,066 species experienced an increase, and 139 species had no stable area across model projections. By IUCN categories, under Criteria A (percent population reduction), 471 species qualify as Critically Endangered (≥ 80% reduction), 688 as Endangered (50-80% reduction), 668 as Vulnerable (30-50% reduction), 1,132 as Near Threatened (0-30% reduction), and 1,075 as Least Concern (≥ 0% increase; Table 2; Figure 2). For the MIROC6 climate model, 2,864 species experienced a reduction of stable area from current suitable area, while 1,163 species experience an increase, and 136 species had no stable area. By IUCN categories, Criteria A, 377 species qualify as Critically Endangered, 544 as Endangered, 622 as Vulnerable, 1,321 as Near Threatened, and 1,170 as Least Concern (Table 2; Figure 2).

**Table 2:** By amphibian order, number of species falling (and percent of total) under IUCN categories (CR – critically endangered; EN – endangered; VU – vulnerable; NT – near threatened; LC – least concern) using Criteria A (precent population reduction) and following the CNRM-CM6-1 and MIROC6 global climate models, based on percent change from current predicted distribution area.

| Order | CNRM-CM6-1 |  |  |  |  | MIROC6 |  |  |  |  |
| --- | --- | --- | --- | --- | --- | --- | --- | --- | --- | --- |
|  | CR | EN | VU | NT | LC | CR | EN | VU | NT | LC |
| Anura | 377<br>(10.8%) | 579<br>(16.6%) | 576<br>(16.5%) | 991<br>(28.4%) | 972<br>(27.8%) | 298<br>(8.5%) | 445<br>(12.7%) | 524<br>(15.0%) | 1171<br>(33.5%) | 1057<br>(30.2%) |
| Caudata | 84<br>(17.4%) | 103<br>(21.4%) | 82<br>(17.0%) | 113<br>(23.4%) | 100<br>(20.7%) | 71<br>(14.7%) | 93<br>(19.3%) | 92<br>(19.1%) | 115<br>(23.9%) | 111<br>(23.0%) |
| Gymnophiona | 10<br>(17.5%) | 6<br>(10.5%) | 10<br>(17.5%) | 28<br>(49.1%) | 3<br>(5.3%) | 8<br>(14.0%) | 6<br>(10.5%) | 6<br>(10.5%) | 35<br>(61.4%) | 2<br>(3.5%) |

**Figure 1:**
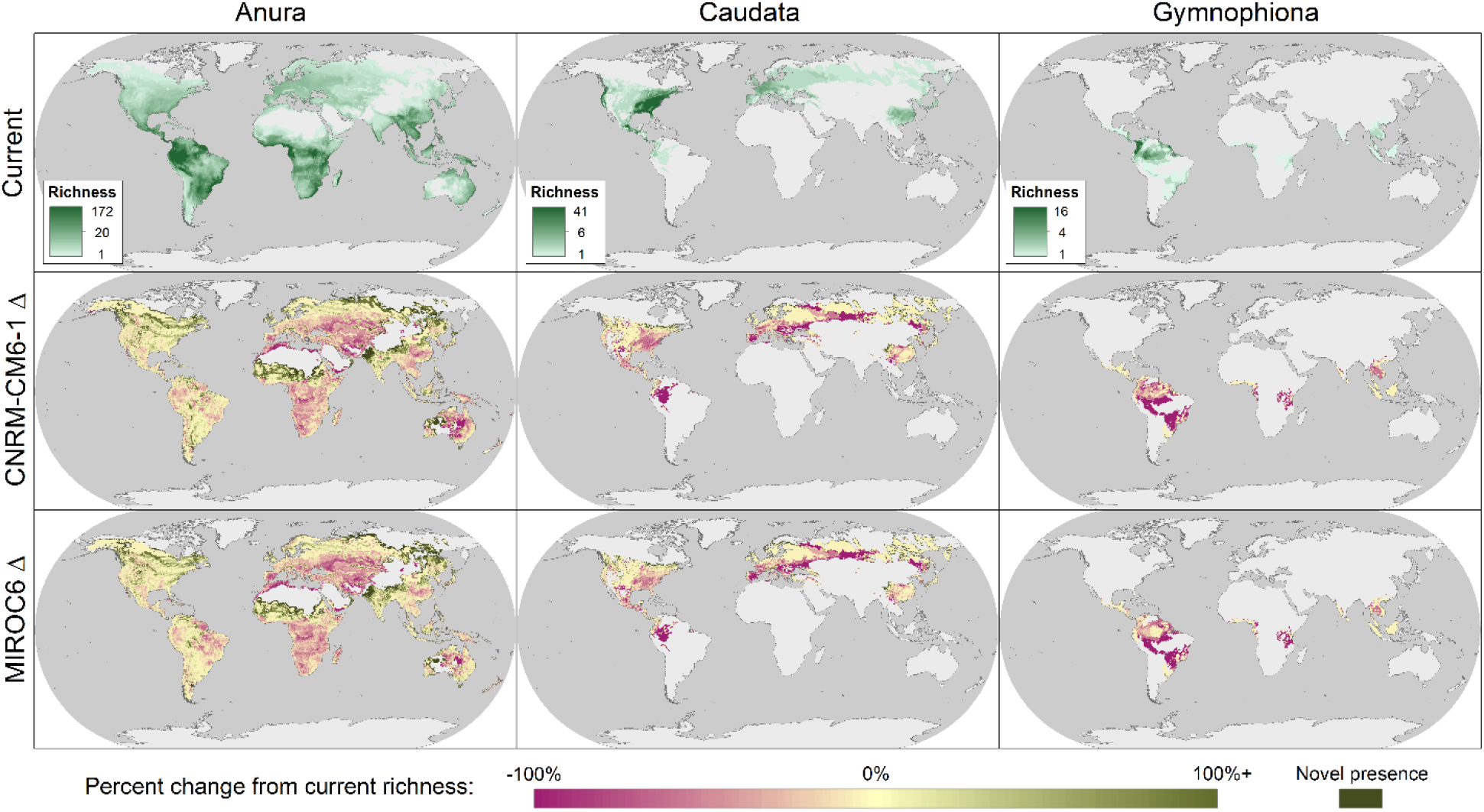
By order, current global richness of amphibians, divided into the three orders, and predicted percent change for refugia under the CNRM-CM6-1 and MIROC6 global climate models.

**Figure 2:**
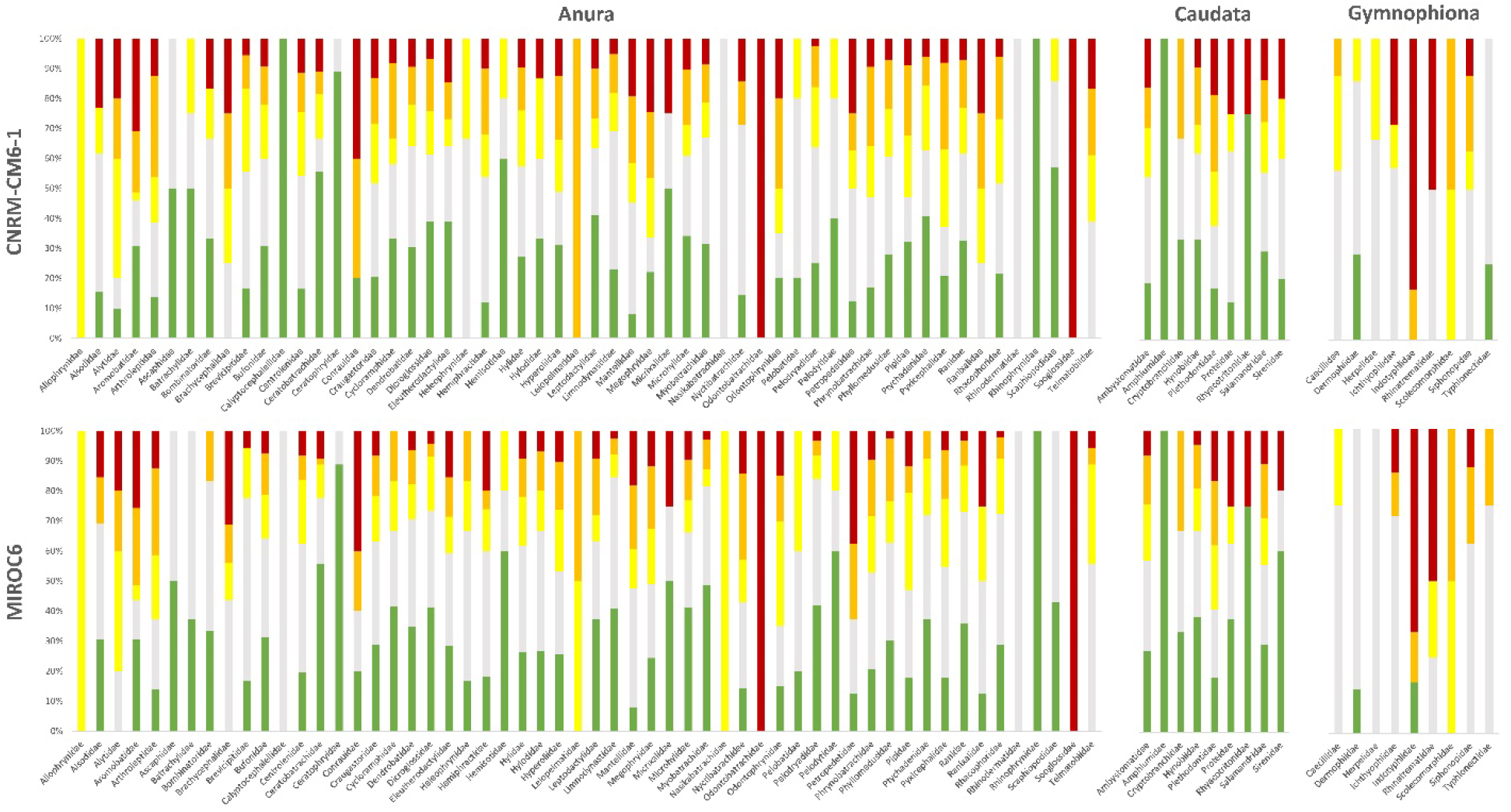
Percent of each amphibian family exhibiting change in refugia area (detailed values in Table S1) from current to MIROC6 and CNRM-CM6-1 global climate models, correlating to IUCN categories using Criteria A (percent population reduction). Categories are as follows: CR – critically endangered (red); EN – endangered (orange); VU – vulnerable (yellow); NT – near threatened (gray); LC – least concern (green).

**Figure 3:**
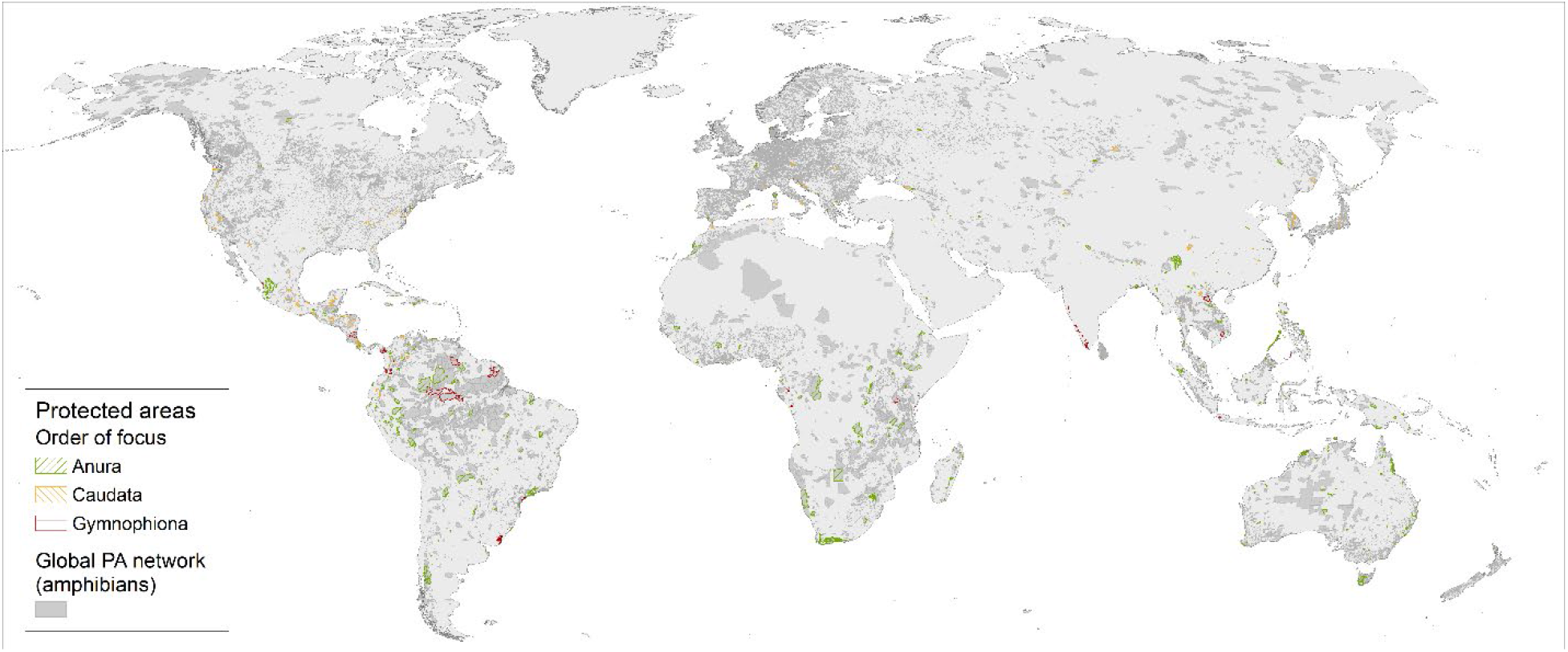
Selected protected areas to maximize number of amphibian species covered in current climate and under CNRM-CM6-1 and MIROC6 climate change models. Protected areas are symbolized by order, with all considered protected areas in gray representing the global protected area network as it applies to amphibians.

### Variables affecting refugia area

When the percent change by family was compared by order (Tukey HSD test) several families emerged as having a significantly greater or lower average change for different climate models. Within Anura, for the CNRM-CM6-1 model, the average percent change of Ceratophrydae species was significantly greater than 16 other families (out of 54), Megophrydae lower than 10 other families, Mantellidae lower than 7 other families, and Arthroleptidae lower than 6 other families. For the MIROC6 model the average percent of change for Ceratobatrachidae was greater than 14 families, Myobatrachidae greater than 10 other families, Ceratophrydae greater than 8 other families, Mantellidae lower than 15 families, and Arthroleptidae less than 9 families. Latitudinal range and absolute value of average latitude had no effect on change in Anura in linear models.

For Caudata, percent change in the CNRM-CM6-1 model was influenced by species’ latitudinal range (F_2,479_ = 2.299, p = 0.0219), with percent change increasing with latitudinal range. In the MIROC6 model, both latitudinal range (F_2,479_ = 2.390, p = 0.0172) and absolute value of latitude (F_2,479_ = 2.143, p = 0.0326) had positive influences on percent change. Family had no effect on percent change within Caudata. Finally, within Gymnophiona, Indotyphlidae had significantly lower change (negative) in CNRM-CM6-1 (p<0.05 for comparison with six out of seven families), with nearly all species having no refugia area. For the MIROC6 model, Indotyphlidae was significantly lower than two of seven families. Latitudinal range and absolute value of average latitude had no effect on change in Gymnophiona in linear models.

### Biomes

Amongst the three orders, specific biomes and regions showed significant differences between current suitable area and refugia depending on the climate change model. In anurans (Table S3), the biomes which had decreased refugia area for both climate change models were evergreen sclerophyllous forests, lake systems, mixed mountain systems, sub-tropical/temperate rain forests/woodlands, tropical dry forests/woodlands, tropical grasslands/savannas, and tropical humid forests. For temperate broad-leaf forests, refugia area was decreased for the CNRM-CM6-1 model, but not the MIROC6 model. In temperate needle-leaf forests/woodlands, refugia area was increased from the current suitable area for both climate models, and tundra communities was increased in the CNRM-CM6-1 model only. In evergreen sclerophyllous forests and mixed mountain systems, refugia in the CNRM-CM6-1 model were significantly smaller than in the MIROC6 model. In tropical dry forests and woodlands, refugia in the MIROC6 model were significantly smaller.

In Caudata (Table S4), the refugia areas were significantly smaller in CNRM-CM6-1 only for cold-winter deserts and sub-tropical and temperate rain forests and woodlands. Refugia areas were significantly smaller for both models in evergreen sclerophyllous forests, mixed mountain systems, temperate broad-leaf forests, temperate grasslands, savannas and warm deserts and semi-deserts. Refugia areas in temperate broad-leaf forests and tropical grasslands and savannas were significantly smaller for the CNRM-CM6-1 model than the MIROC6 model. Finally, Gymnophiona (Table S5) had significantly smaller refugia areas in mixed mountain systems for both models and in tropical dry forests/woodlands and tropical humid forests for only the MIROC6 model.

### Regions

Where they are currently present, anurans (Table S6) will experience decreased refugia area in 10 of 24 global regions for CNRM-CM6-1 and 11 global regions for MIROC6. Refugia area will increase in five regions for CNRM-CM6-1 and three regions for MIROC6, mostly in higher latitudes in the northern hemisphere. Caudata (Table S7) climate refugia will decrease in eight of 15 regions for CNRM-CM6-1 and five regions for MIROC6. Finally, Gymnophiona (Table S8) refugia areas will decrease in one of five tested regions for CNRM-CM6-1 and 2 for MIROC6.

## Discussion

While the outcomes of the CNRM-CM6-1 and MIROC6 models were similar, the MIROC6 climate model resulted in less negative impact overall. This fits with the MIROC6 model having a lower climate sensitivity. Our results indicate that the predicted percent change for refugia is generally negative for all orders, with Caudata and Gymnophiona showing the highest declines (−34.1% and -37.5% for CNRM-CM6-1, respectively; -29.8% and -32.6% for MIROC6, respectively). Interestingly, current diversity hotspots were not necessarily impacted the most; more marginal areas may experience a greater decline in terms of percent species losing refugia area. Furthermore, the number of species falling under IUCN categories CR, EN, and VU is expected to increase under both climate models, indicating a high risk of extinction for specific amphibian species. We also found that a total of 139 and 136 species will have no climate refugia under CNRM-CM6-1 and MIROC6 climate models, respectively. All species with no future climate refugia had current predicted suitable area less than 0.4 km^2^, further highlighting the negative impact of climate change on rare amphibians with geographically narrow populations.

Among the families that will be less negatively affected by climate change are Ceratophryidae (horned frogs), Ceratobatrachidae (ground frogs) and Myobatrachidae (Australian ground and water frogs). Species in these families all have, to some extent, adaptations to desiccation. Although Ceratophryidae have been predicted to be negatively associated with climate change in PAs (Loyola et al., 2014), their larval morphology (Faivovich et al., 2014) and rapid larval development (Fabrezi, 2011; Quinzio et al., 2006) relate to adaption to semi-arid environments. Additionally, adults are known to burrow into humid soil in times of low humidity and even produce a cocoon that reduces water loss during dormancy (Bastos and Abe, 1998; McClanahan et al., 1976). Ceratobatrachidae have direct development (Narayan et al., 2011), or no tadpole stage, which generally relates to less reliance on permanent or even ephemeral bodies of water. Some species in the family utilize phytotelmata, or pockets of water tapped by plant leaves, to breed (Bucol et al., 2019). Finally, among Myobatrachidae, *Arenophryne rotunda* is a sand burrowing species inhabiting arid and semi-arid environments (Cartledge et al., 2006), and *Pseudophryne australis* shows asynchrony of metamorphosis indicating both plasticity and bet-hedging strategies (Thumm and Mahony, 2006). In any case, this family has wide-ranging development strategies even within genera (Anstis, 2010; Knowles et al., 2014), indicating a potential for high levels of plasticity within the family.

Families that will be more negatively impacted are concentrated in specific geographic regions or have specialized or narrow niches. Arthroleptidae, which are native to sub-saharan Africa (Blackburn, 2008; Portillo and Greenbaum, 2014), usually live in leaf litter and may be more susceptible to environmental changes due to high levels of specialization. Mantellidae is a family restricted to Madagascar and Mayotte (Andreone et al., 2002). Being island locked means there is no way for species to migrate to suitable climates if the islands become inhospitable. This family includes specialization in the form of consuming ants and mites for dietary alkaloids to produce toxins in the skin (Woodhead et al., 2007). For Megophryidae, it is unclear what traits may increase their sensitivity to climate change, but some species are found at high elevations (Rowley et al., 2013; Subba et al., 2015) which will undoubtedly experience the effects of climate change. Indotyphlidae, a family of small caecilians, has a patchy distribution in Africa, India and the Seychelles (Wilkinson et al., 2011). Species in this family may be more negatively impacted because of the patchy and restricted distributions, island distributions, and distributions in regions that will likely experience greater drought impacts of climate change.

Although this type of climate change modeling provides valuable insights into global patterns of impacts on amphibians, it is important to note the limitations. These include consideration of habitats and microhabitats, for example vernal pools (Cartwright et al., 2022), that are not possible to model on a global scale as well as species with too few occurrence records to model. However, in the case of the latter, an assumption can be made that any refugia area for such species will be small and likely overlapping with larger biodiversity patterns, in which case it is even more important to ensure protection of such areas of high species richness. Additionally, such modeling does not take into account the potential for adaptation to changing climates. In the case of some well-studied species, trait-based modeling can be implemented (Benito Garzón et al., 2019), but this is dependent on knowledge of species’ phenotypic plasticity. Another method that may be useful for large-scale modeling is latitudinal adjustment of bioclimatic variables used to build models, which has already been shown to improve model accuracy in both narrow and wide-ranging species (Andersen et al., 2022).

This study highlights the importance of identifying areas of refugia, or areas of high climatic stability, to protect the survival of amphibian species in the face of rapid climate change. By identifying a subset of protected areas to conserve the maximum number of species both currently and in different climate change scenarios, we provide managers of these protected areas a focus to conserve amphibian richness to strengthen the larger network of protected areas. These areas will be critical for the survival of species and managers can help develop effective conservation strategies that protect not only individual species but also the ecosystems and services they provide. The databases of species’ current and refugia areas extracted to geographic regions, biomes, global protected areas, and key biodiversity areas (KBAs) can serve as a tool for policymakers, researchers, and conservation practitioners to identify critical areas for conservation.

## Notes

### Competing Interest Statement

The authors have declared no competing interest.

