## Supplementary material for "A slippery slope: assessing the amphibian extinction crisis through the lens of climate refugia": Tables S3-8

Table S3: Udvardy biome difference in refugia areas for Anura. Significant differences between current area and CNRM-CM6-1 and MIROC6 refugia areas are indicated in bold. Negatives in the CNRM-Current and MIROC-Current columns indicate decreased climate change refugia area, while negatives in the CNRM-MIROC column indicates CNRM has a smaller area of refugia than MIROC. Area is measured in number of cells of 0.04*0.04 decimal degrees, or ~3.7*3.7 km.

|  | Difference | | | P(T<=t) two-tail | | |
| --- | --- | --- | --- | --- | --- | --- |
|  | CNRM-Current | MIROC-Current | CNRM-MIROC | CNRM-Current | MIROC-Current | CNRM-MIROC |
| Cold-winter deserts | -142.2 | -356.2 | 214 | 0.7198 | 0.4117 | 0.2404 |
| Evergreen Sclerophyllous forests | **-796.8** | **-570.2** | **-226.6** | **<0.0001** | **0.0019** | **0.0065** |
| Lake systems | **-180.3** | **-219.0** | 38.7 | **0.0270** | **0.0015** | 0.2615 |
| Mixed island systems | -122.2 | -44.6 | -77.6 | 0.2401 | 0.5856 | 0.0809 |
| Mixed mountain systems | **-425.7** | **-333.3** | **-92.4** | **<0.0001** | **<0.0001** | **0.0042** |
| Sub-tropical / Temperate rain forests / Woodlands | **-1492.2** | **-1402.4** | -89.8 | **<0.0001** | **<0.0001** | 0.3750 |
| Temperate broad-leaf forests | **-2031.4** | -1631.0 | -400.4 | **0.0284** | 0.0611 | 0.1723 |
| Temperate grasslands | -229.6 | -643.4 | 413.8 | 0.5917 | 0.1502 | 0.0542 |
| Temperate needle-leaf forests / Woodlands | **17760.2** | **15651.9** | 2108.2 | **<0.0001** | **<0.0001** | 0.3986 |
| Tropical dry forests / Woodlands | **-924.0** | **-1136.0** | **212.0** | **<0.0001** | **<0.0001** | **0.0137** |
| Tropical grasslands / Savannas | **-773.9** | **-911.9** | 138.0 | **0.0093** | **0.0004** | 0.2894 |
| Tropical humid forests | **-1942.8** | **-1861.2** | -81.7 | **<0.0001** | **<0.0001** | 0.2754 |
| Tundra communities | **9511.4** | 7290.3 | 2221.1 | **0.0350** | 0.0896 | 0.5710 |
| Warm deserts / semi-deserts | -171.8 | -232.5 | 60.7 | 0.4509 | 0.1748 | 0.4701 |

Table S4: Udvardy biome difference in refugia areas for Caudata. Significant differences between current area and CNRM-CM6-1 and MIROC6 refugia areas are indicated in bold. Negatives in the CNRM-Current and MIROC-Current columns indicate decreased climate change refugia area, while negatives in the CNRM-MIROC column indicates CNRM has a smaller area of refugia than MIROC. Area is measured in number of cells of 0.04*0.04 decimal degrees, or ~3.7*3.7 km.

|  | Difference | | | P(T<=t) two-tail | | |
| --- | --- | --- | --- | --- | --- | --- |
|  | CNRM-Current | MIROC-Current | CNRM-MIROC | CNRM-Current | MIROC-Current | CNRM-MIROC |
| Cold-winter deserts | **-313.8** | -224.5 | -89.2 | **0.0491** | 0.1213 | 0.0887 |
| Evergreen Sclerophyllous forests | **-883.2** | **-912.6** | 29.4 | **0.0010** | **0.0008** | 0.4558 |
| Lake systems | -299.6 | -235.7 | -63.9 | 0.1914 | 0.3712 | 0.6172 |
| Mixed island systems | -29.3 | -5.0 | -24.3 | 0.3716 | 0.6159 | 0.2942 |
| Mixed mountain systems | **-359.9** | **-297.2** | -62.7 | **0.0002** | **0.0025** | 0.1319 |
| Sub-tropical / Temperate rain forests / Woodlands | **-382.7** | -251.8 | -130.9 | **0.0177** | 0.0559 | 0.0944 |
| Temperate broad-leaf forests | **-6071.1** | **-4882.0** | **-1189.2** | **<0.0001** | **<0.0001** | **<0.0001** |
| Temperate grasslands | **-1281.3** | **-1365.9** | 84.6 | **0.0058** | **0.0042** | 0.4158 |
| Temperate needle-leaf forests / Woodlands | -1919.3 | -1663.1 | -256.3 | 0.7418 | 0.7738 | 0.3438 |
| Tropical dry forests / Woodlands | **-120.9** | **-101.1** | -19.8 | **<0.0001** | **<0.0001** | 0.1913 |
| Tropical grasslands / Savannas | -471.3 | -297.8 | **-173.5** | 0.2139 | 0.3189 | **0.0489** |
| Tropical humid forests | -678.7 | -638.9 | -39.8 | 0.1883 | 0.1790 | 0.3527 |
| Warm deserts / semi-deserts | **-591.4** | **-598.4** | 7.0 | **0.0412** | **0.0256** | 0.8995 |

Table S5: Udvardy biome difference in refugia areas for Gymnophiona. Significant differences between current area and CNRM-CM6-1 and MIROC6 refugia areas are indicated in bold. Negatives in the CNRM-Current and MIROC-Current columns indicate decreased climate change refugia area, while negatives in the CNRM-MIROC column indicates CNRM has a smaller area of refugia than MIROC. Area is measured in number of cells of 0.04*0.04 decimal degrees, or ~3.7*3.7 km.

|  | Difference | | | P(T<=t) two-tail | | |
| --- | --- | --- | --- | --- | --- | --- |
|  | CNRM-Current | MIROC-Current | CNRM-MIROC | CNRM-Current | MIROC-Current | CNRM-MIROC |
| Mixed mountain systems | **-1219.5** | **-851.2** | **-368.3** | **<0.0001** | **<0.0001** | **0.0863** |
| Sub-tropical / Temperate rain forests / Woodlands | -10723.5 | -10059.8 | -663.8 | 0.3598 | 0.3714 | 0.1692 |
| Tropical dry forests / Woodlands | -2042.7 | **-2042.6** | -0.1 | 0.0517 | **0.0275** | 0.9998 |
| Tropical grasslands / Savannas | -1967.1 | -1559.7 | -407.4 | 0.0961 | 0.0628 | 0.4273 |
| Warm deserts / semi-deserts | -971.8 | -1023.8 | 52.0 | 0.2239 | 0.2207 | 0.2862 |

Table S6: World region difference in refugia areas for Anura. Significant differences between current area and CNRM-CM6-1 and MIROC6 refugia areas are indicated in bold. Negatives in the CNRM-Current and MIROC-Current columns indicate decreased climate change refugia area, while negatives in the CNRM-MIROC column indicates CNRM has a smaller area of refugia than MIROC. Area is measured in number of cells of 0.04*0.04 decimal degrees, or ~3.7*3.7 km.

|  | Difference | | | P(T<=t) two-tail | | |
| --- | --- | --- | --- | --- | --- | --- |
|  | CNRM-Current | MIROC-Current | CNRM-MIROC | CNRM-Current | MIROC-Current | CNRM-MIROC |
| Asiatic Russia | **14199.0** | **11537.4** | 2661.6 | **0.0084** | **0.0166** | 0.5304 |
| Australia/New Zealand | **-1961.8** | -341.2 | **-1620.5** | **0.0014** | 0.3249 | **<0.0001** |
| Caribbean | -2.7 | **-103.5** | **100.8** | 0.9626 | **0.0133** | **0.0169** |
| Central America | **-421.1** | **-449.8** | 28.8 | **0.0009** | **<0.0001** | 0.5786 |
| Central Asia | -1839.1 | -3985.2 | 2146.1 | 0.4858 | 0.2409 | 0.1330 |
| Eastern Africa | **-2619.5** | **-3367.5** | **748.0** | **<0.0001** | **<0.0001** | **<0.0001** |
| Eastern Asia | **-1496.2** | **-782.0** | **-714.2** | **0.0001** | **0.0130** | **<0.0001** |
| Eastern Europe | -7953.4 | **-9744.0** | 1790.6 | 0.0922 | **0.0309** | 0.3311 |
| European Russia | -1644.2 | -5231.0 | 3586.8 | 0.7300 | 0.2439 | 0.1339 |
| Melanesia | **-392.9** | **-191.4** | **-203.5** | **<0.0001** | **0.0017** | **<0.0001** |
| Micronesia | **12.3** | 11.0 | 1.3 | **0.0483** | 0.0908 | 0.3632 |
| Middle Africa | **-5816.8** | **-5883.2** | 66.4 | **<0.0001** | **<0.0001** | 0.8297 |
| Northern Africa | -403.1 | -801.3 | 398.3 | 0.5581 | 0.1717 | 0.0597 |
| Northern America | **2980.8** | **3271.4** | -290.6 | **0.0170** | **0.0014** | 0.5635 |
| Northern Europe | **7094.1** | **6322.8** | 771.3 | **0.0003** | **0.0002** | 0.5451 |
| Polynesia | -117.3 | -156.3 | 39.0 | 0.1513 | 0.0830 | 0.3426 |
| South America | **-2544.2** | **-2809.5** | 265.3 | **<0.0001** | **<0.0001** | 0.1190 |
| Southeastern Asia | **-1003.7** | **-376.0** | **-627.7** | **<0.0001** | **0.0180** | **<0.0001** |
| Southern Africa | **-2385.9** | **-2095.0** | -291.0 | **0.0003** | **<0.0001** | 0.2065 |
| Southern Asia | -412.3 | -23.1 | **-389.2** | 0.0840 | 0.8952 | **0.0443** |
| Southern Europe | **-1807.1** | **-2868.0** | **1060.9** | **0.0077** | **0.0005** | **0.0062** |
| Western Africa | **2112.2** | 1240.1 | **872.1** | **0.0091** | 0.0678 | **<0.0001** |
| Western Asia | 301.8 | 91.9 | 209.9 | 0.3746 | 0.8100 | 0.5577 |
| Western Europe | -900.5 | -1314.9 | 414.3 | 0.5239 | 0.4031 | 0.2668 |

Table S7: World region difference in refugia areas for Caudata. Significant differences between current area and CNRM-CM6-1 and MIROC6 refugia areas are indicated in bold. Negatives in the CNRM-Current and MIROC-Current columns indicate decreased climate change refugia area, while negatives in the CNRM-MIROC column indicates CNRM has a smaller area of refugia than MIROC. Area is measured in number of cells of 0.04*0.04 decimal degrees, or ~3.7*3.7 km.

|  | Difference | | | P(T<=t) two-tail | | |
| --- | --- | --- | --- | --- | --- | --- |
|  | CNRM-Current | MIROC-Current | CNRM-MIROC | CNRM-Current | MIROC-Current | CNRM-MIROC |
| Asiatic Russia | -7048.6 | -7733.2 | 684.6 | 0.5201 | 0.5082 | 0.3620 |
| Central America | **-487.7** | **-510.3** | 22.5 | **<0.0001** | **<0.0001** | 0.5236 |
| Eastern Asia | **-1978.1** | **-1696.5** | -281.6 | **0.0084** | **0.0122** | 0.1401 |
| Eastern Europe | **-9418.2** | -9222.2 | -196.0 | **0.0431** | 0.0694 | 0.8589 |
| European Russia | -10043.5 | -10720.5 | 677.0 | 0.2726 | 0.2255 | 0.5430 |
| Northern Africa | -498.7 | -479.3 | -19.3 | 0.1933 | 0.1658 | 0.6756 |
| Northern America | **-4147.0** | **-3025.7** | **-1121.3** | **<0.0001** | **<0.0001** | **<0.0001** |
| Northern Europe | 4256.6 | 4561.2 | -304.6 | 0.1212 | 0.1097 | 0.0582 |
| South America | -3074.4 | -2754.6 | -319.8 | 0.2217 | 0.2383 | 0.0914 |
| Southeastern Asia | -591.6 | -420.3 | -171.3 | 0.1292 | 0.0781 | 0.2856 |
| Southern Asia | **-254.8** | -191.6 | -63.2 | **0.0389** | 0.0560 | 0.4599 |
| Southern Europe | **-2681.1** | **-2871.1** | 190.0 | **0.0026** | **0.0019** | 0.1640 |
| Western Asia | **-604.4** | -327.1 | **-277.4** | **0.0047** | 0.1393 | **0.0223** |
| Western Europe | **-4752.4** | **-4669.7** | -82.7 | **0.0108** | **0.0131** | 0.8632 |

Table S8: World region difference in refugia areas for Gymnophiona. Significant differences between current area and CNRM-CM6-1 and MIROC6 refugia areas are indicated in bold. Negatives in the CNRM-Current and MIROC-Current columns indicate decreased climate change refugia area, while negatives in the CNRM-MIROC column indicates CNRM has a smaller area of refugia than MIROC. Area is measured in number of cells of 0.04*0.04 decimal degrees, or ~3.7*3.7 km.

|  | Difference | | | P(T<=t) two-tail | | |
| --- | --- | --- | --- | --- | --- | --- |
|  | CNRM-Current | MIROC-Current | CNRM-MIROC | CNRM-Current | MIROC-Current | CNRM-MIROC |
| Central America | -265.5 | **-511.1** | 245.6 | 0.0781 | **0.0200** | 0.2147 |
| Eastern Africa | -4337.3 | -4564.6 | 227.3 | 0.3026 | 0.2892 | 0.2152 |
| Middle Africa | -3318.5 | -3368.5 | 50.0 | 0.0949 | 0.0613 | 0.9431 |
| South America | **-8694.2** | **-7278.0** | -1416.1 | **0.0276** | **0.0183** | 0.2527 |
| Southeastern Asia | -5728.8 | -4079.2 | -1649.7 | 0.3113 | 0.2994 | 0.3427 |
